# Antigen flexibility supports the avidity of hemagglutinin-specific antibodies at low antigen densities

**DOI:** 10.1101/2024.11.20.624560

**Authors:** Ananya Benegal, Yuanyuan He, Katilyn Ho, Giselle Groff, Zijian Guo, Michael D. Vahey

## Abstract

The receptor-binding protein of influenza A virus, hemagglutinin (HA), is the most abundant protein on the viral surface. While high densities of HA are thought to improve cellular attachment by increasing avidity for the viral receptor, they may also increase the avidity of neutralizing antibodies. The tradeoff between these two competing effects of avidity is not well understood. To better understand how features of the viral surface influence antibody avidity, we developed fluorescence-based assays to measure dissociation kinetics and steady-state binding of antibodies to intact virions. Focusing on two antibodies that bind to the HA head domain (S139/1 and C05), we confirm that binding orientations that favor bivalent attachment of antibodies to the viral surface can offset weak monovalent affinity by facilitating crosslinking. By modulating HA density in both engineered viruses and synthetic nanoparticles, we find that bivalent antibody binding remains resilient down to one-tenth the HA density on the viral surface and, in the case of C05, that antibody occupancy increases at these lowest densities. Finally, using a combination of structure-guided modeling and antibodies that lock HA in a tilted conformation, we identify flexibility of the HA ectodomain as an additional determinant of antibody avidity. Together, these results establish features of the viral surface that help support or suppress the binding of neutralizing antibodies.

## INTRODUCTION

Influenza A viruses (IAVs) cause seasonal epidemics and occasional pandemics with global mortality in the hundreds of thousands to millions each year^1^. Influenza virions are covered with a dense arrangement of the spike proteins hemagglutinin (HA) and neuraminidase (NA). HA, the receptor binding protein, is the most abundant protein on the IAV surface, with hundreds to thousands of copies per virion. The monovalent binding affinity between HA and the viral receptor, sialic acid, is very low; as a result, only through multiple simultaneous interactions can a virion stably attach to a host cell^2,3^. The high surface density of HA is thought to play a critical role in facilitating viral binding to the host cell surface during infection by leveraging high binding avidity to compensate for low affinity^4–6^ (Fig. 1A).

**Figure 1:**
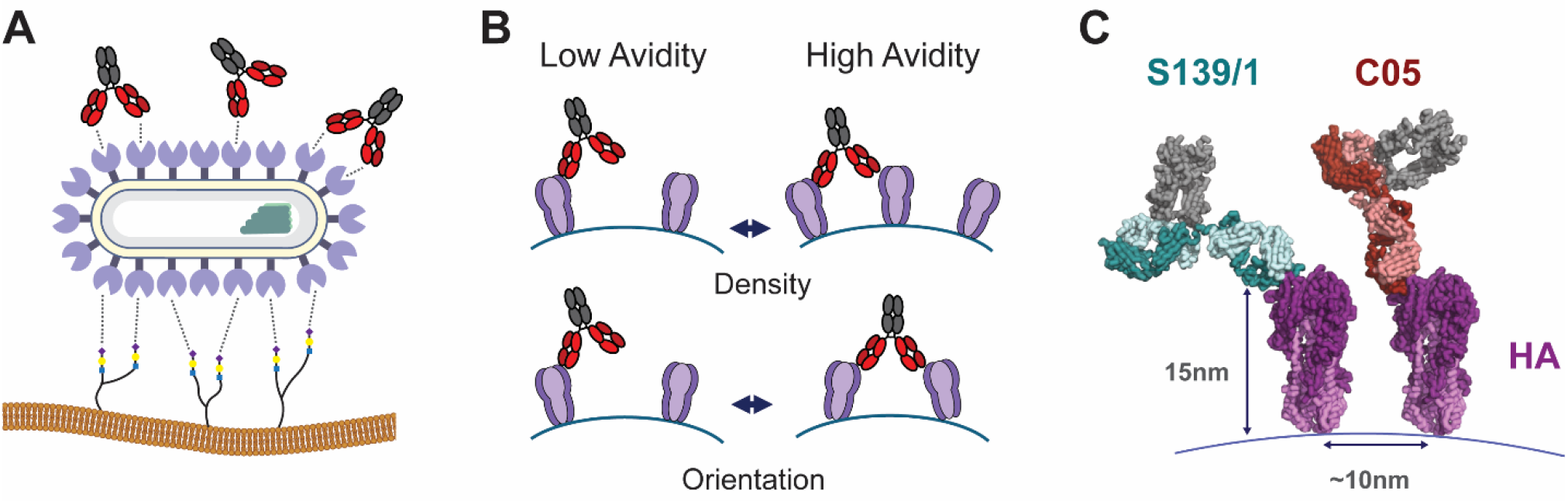
High densities of HA on the IAV surface can contribute to host cell attachment and the binding of neutralizing antibodies. (A) HAs on the viral surface bind to multiple sialylated receptors (depicted in the lower part of the image), increasing receptor-binding avidity. High densities of HA also present opportunities for bivalent attachment of antibodies (upper part of the image). (B) Factors that may influence the ability of antibodies to bind bivalently are the density of HA on the virion surface and the degree to which they can tilt and rotate. (C) Structural model of S139/1 (PDB ID 4GMS) and C05 (4FQR) bound to the HA head domain (3LZG). Distances indicate the height of the HA ectodomain (∼15nm) and typical nearest-neighbor spacing within the viral membrane (∼10nm).

Avidity can also play an antiviral role, by contributing to the binding of neutralizing antibodies. Antibodies against HA are among the most effective protection against influenza virus infection, blocking viral attachment, entry, or release, and engaging with immune cell effector functions^7^. Since natural antibodies contain at least two antigen binding fragments (Fabs), the apparent binding affinity of an antibody can be enhanced through avidity, in some cases by multiple orders of magnitude^8–10^. While this effect is widely recognized, the extent to which specific antibodies leverage avidity is shaped by multiple factors and is therefore difficult to predict. These factors could include the antibody isotype, as well as the location, density, and organization of target epitopes on the antigenic surface (Fig. 1B). For example, broadly neutralizing antibodies against HA often bind to the more conserved membrane-proximal stalk^11–13^, which is less accessible for bivalent binding^14–16^. In contrast, head-binding antibodies target epitopes that are more sterically accessible, which may be more conducive to bivalent binding^8,9,17^. Understanding how the features of an antigenic surface enhance or suppress antibody avidity could inform the design of next-generation vaccines^18,19^.

To investigate how the features of antigenic surfaces influence antibody avidity, we developed fluorescence-based assays for measuring antibody binding kinetics to intact virions. Using this approach, we determined antigen cross-linking rates to compare the extent to which specific antibodies leverage bivalent binding. Through comparisons of two HA head-binding antibodies – S139/1^8^ and C05^9^ (Fig. 1C) – we confirmed that a favorable binding orientation that promotes crosslinking can compensate for weak monovalent affinity. Using these antibodies as models, we tested the effect of decreasing antigen density using both virions that contain a fluorescent decoy HA as well as synthetic nanoparticles. We find that both antibodies are surprisingly resilient to changes in surface HA density. This resilience depends in part on the ability of HA to tilt extensively about its membrane anchor. Collectively, these findings present a framework for dissecting antibody avidity, and suggest that antigen flexibility – in addition to antigen surface density – may play an outsized role.

## RESULTS

### A fluorescence-based assay to capture antibody crosslinking kinetics

To determine crosslinking rates of HA-specific antibodies, we used a fluorescence microscopy-based approach to measure antibody dissociation kinetics to intact virus particles. Imaging antibody binding to virions allows us to characterize binding kinetics under conditions where Has are displayed in a physiologically accurate way. Here, we focus on S139/1^8^ and C05^9^, two head-binding antibodies that interact with HA with low monovalent affinity as Fabs but high avidity as bivalent IgG1. Both experimental and computational studies have shown that IgG antibodies have considerable flexibility in their hinge region that allow the antibody to sample conformations that promote bivalent binding^20–23^. We model the dissociation of bivalent antibodies as a combination of two rates: 1) the off-rate of a single Fab arm, and 2) the crosslinking rate (‘*k*_*x*_’) that describes the binding of a free HA by a singly-bound IgG (Fig. 2A). This crosslinking rate will depend on the association rate of the HA-Fab interaction, the steric accessibility of the epitope, the geometry of binding, and the density of available HAs. To measure these rates, whole virions are immobilized in a flow chamber, and fluorescently labeled antibody is introduced and allowed to bind to equilibrium. The antibody is then washed out, and dissociation is measured by loss of fluorescence signal over time (Fig. 2B). To confirm that the loss in signal is not a result of photobleaching, we image antibody bound at steady-state and do not observe a decrease in fluorescence (Fig. S1A). The initial dissociation rate of the antibody provides a value for *k*_*off*_ which we can compare between different antibodies in both IgG1 and Fab formats. By using a continuous-time model in which each arm of an IgG is assumed to dissociate at the rate of the corresponding Fab, we fit the experimentally-determined *k*_*off*_ values of the Fab and IgG pairs to calculate *k*_*x*_ (Fig. S1B, details provided in Methods).

**Figure 2:**
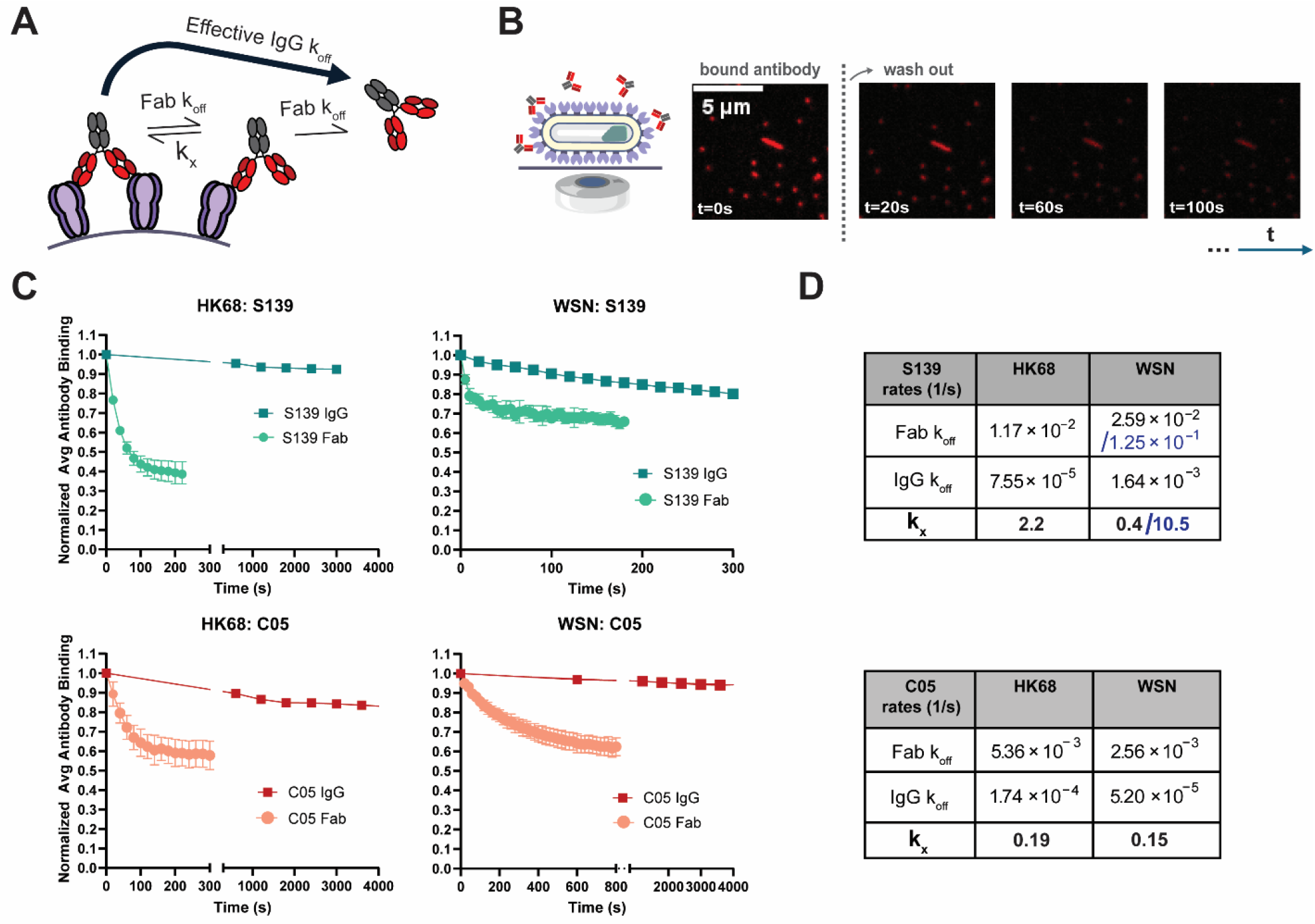
S139/1 and C05 IgGs are enhanced by avidity to different extents. (A) The effective off-rate of an IgG antibody (IgG k_off_) is modeled by the combination of two rates: the off-rate for each of the Fab arms (Fab *k*_*off*_), and the cross-linking rate (*k*_*x*_) at which the second Fab binds when the first Fab is bound. (B) Schematic and images from the assay to measure dissociation kinetics. IAV immobilized onto a flowchamber surface is imaged after equilibrated fluorescent antibody is rapidly washed away. (C) Normalized Fab and IgG dissociation curves for both S139/1 and C05 against the HK68 and WSN strains. Data are combined from three biological replicates at two antibody concentrations each and normalized to the fluorescence intensity at *t*=0 for each sample. Time between acquisition is set to 5, 20, or 600 seconds depending on relative rate of dissociation. (D) Kinetic parameters for S139/1 and C05 Fab/IgG against WSN33 and HK68. The dissociation rates for each antibody/strain combination are determined from the initial rates of fluorescence loss. The crosslinking rate *k*_*x*_ is fit for each pair by simulating the effective IgG *k*_*off*_ for a given Fab *k*_*off*_ for a range of *k*_*x*_ values. A *k*_*x*_ value estimated from data in Lee *et al*. is shown in blue for S139/1 Fab against WSN33.

### Antigen crosslinking can slow antibody dissociation by as much as two orders of magnitude

We measured IgG and Fab dissociation kinetics for S139/1 and C05 against the H1N1 strain A/WSN/1933 (WSN33) and the H3N2 strain A/Hong Kong/1/1968 (HK68) (Fig. 2C). Because the rapid dissociation of the S139/1 Fab against WSN33 HA approaches the sampling frequency of our assay, our measured *k*_*off*_ in this case likely underestimates the true dissociation rate; we therefore also compared an off-rate estimated from reported BLI data^8^. We find that for both antibodies, the crosslinking rate is faster than the monovalent off-rate by at least an order of magnitude, and thus significantly boosts the ability of the IgG form of the antibody to remain bound (Fig. 2D). In general, avidity will enhance affinity when new crosslinks form at a timescale similar to or faster than the monovalent dissociation rate (Fig. S1C). Although antibody loading at very high concentrations could favor monovalent over bivalent attachment and lead to initial rapid dissociation kinetics, our measured dissociation rates are insensitive to the loading concentration of the antibody within the range that we test (Fig. S1D & E). Avidity of these antibodies results in a decrease in dissociation rate of up to 150-fold, for S139/1 against HK68 HA. Although the S139/1 Fab dissociates twice as fast from HK68 HA as the C05 Fab, it’s higher rate of crosslinking leads to slower dissociation when the two antibodies are compared as IgGs. This enhancement in effective affinity by multiple orders of magnitude parallels previous measurements for these and other antibodies^10,17^.

### Decreasing HA surface density 25% through introduction of an HA decoy does not affect antibody binding avidity

The ability of an antibody to bind bivalently to an antigenic surface will depend on the availability of antigens that are suitably positioned, and should be reduced if HA density is sufficiently low. To experimentally reduce the HA surface density, we expressed a fluorescent HA decoy that competes with wildtype HA for packaging into virions in infected cells (Fig. 3A). The decoy is comprised of the cytoplasmic tail and transmembrane domain of H1 HA, while the HA ectodomain is replaced by the GCN4-pII trimerization domain^24^ fused to the fluorescent protein super-ecliptic pHluorin (SEP)^25^. To prevent co-trimerization of the wildtype HA and the decoy (‘SEP-HA’) through interactions between their transmembrane domains, we found that it is important that the SEP-HA transmembrane domain be derived from a different subtype than the virus used for infection; thus, for these experiments, we created an MDCK stable cell line expressing SEP-HA derived from WSN33 and infected with HK68 virus. This approach results in viruses that are morphologically similar to wildtype, albeit with a higher number of total particles released over the course of infection (Fig. S2 A-C). Virus collected from these infections is then used to determine the reduction of HA density that occurs due to competition between wildtype and SEP-HA, and the corresponding effect on antibody binding. Fluorescently labeled antibody is added to immobilized virus and allowed to bind to equilibrium, and the Fab and IgG steady-state binding is compared for the wild-type and mutant virions.

**Figure 3.**
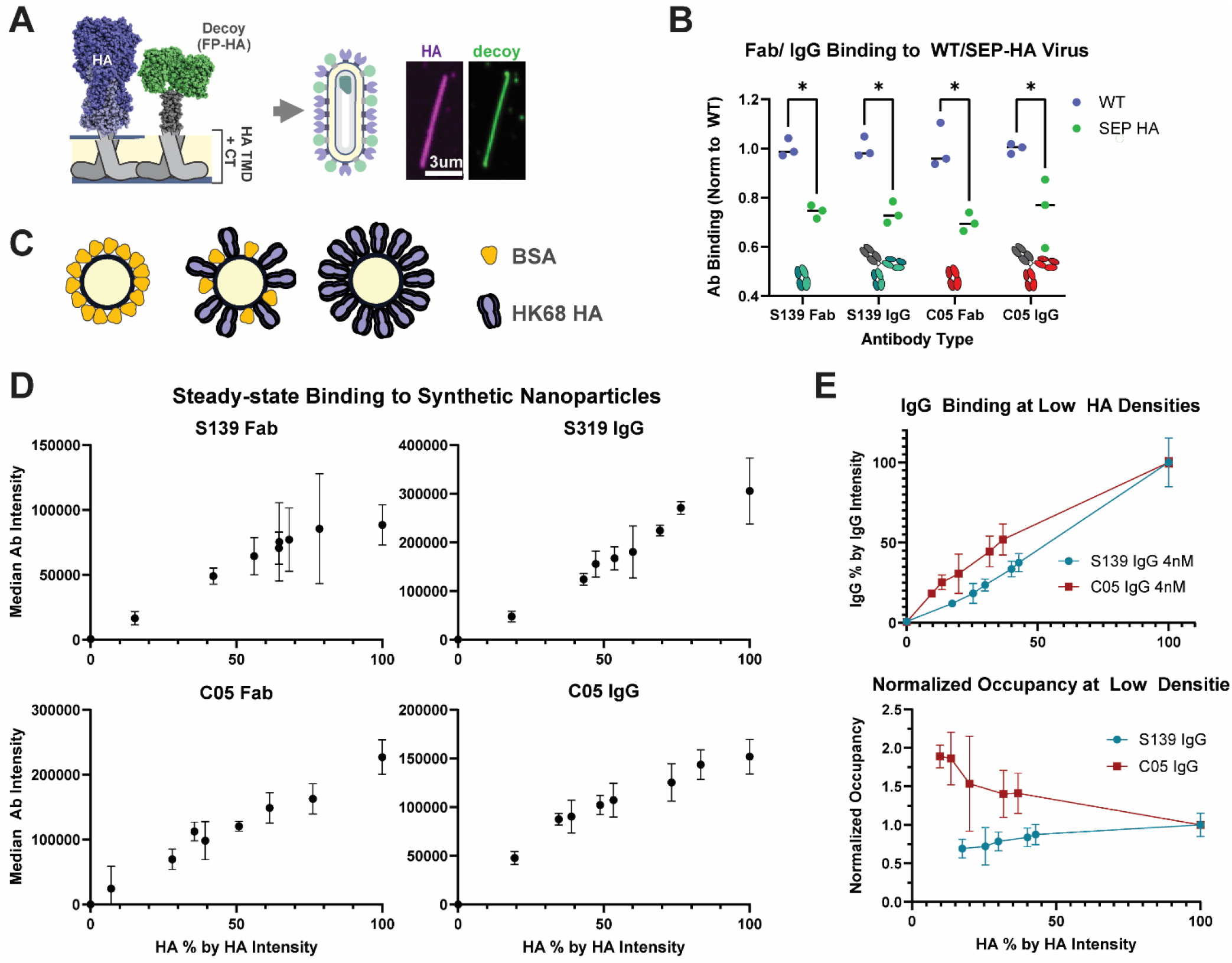
S139/1 and C05 avidity persists at ten-fold reductions in HA density. (A) Schematic and representative image of the fluorescent HA decoy used to reduce the HA density on the viral surface. The decoy (SEP-HA) contains the cytoplasmic tail and transmembrane domain of WSN33 HA, while the head domain is replaced by super-ecliptic pHluorin. (B) Steady-state binding of the antibodies to the WT and SEP-HA versions of the virus measured by fluorescence intensity. Quantification is performed on large filamentous virions to ensure high signal-to-noise. Measured intensities are normalized to the mean of the WT condition for each pair. P-values are determined by multiple unpaired t-tests; significance is indicated by a p-value below 0.05. (C) Schematic of the experimental approach to decrease HA density using synthetic nanoparticles. C-terminally biotinylated HK68 HA ectodomain is bound to streptavidin-coated spheres and diluted with varying concentrations of biotinylated BSA to titrate the relative HA density. (D) Steady-state binding of Fab and IgG antibodies as a function of relative HA density. Relative HA density is determined by normalizing mean HA signal intensity to the 100% HA condition in each replicate. Three separate preparations of beads are measured for each condition. (E) Comparison of S139/1 and C05 IgG binding at lower HA densities. Top: both HA and the antibody fluorescence intensities are also normalized to the mean 100% HA condition for each antibody for all three replicates. Bottom: antibody occupancy as a function of HA density. For each point, %IgG from the top plot is divided by the corresponding %HA.

We reasoned that the monovalent binding of a Fab to an epitope at the apex of HA should be unaffected by changes in HA density. Accordingly, we used a C05 Fab fragment as a proxy for HA abundance to determine the average extent to which wildtype HA is diluted by the SEP-HA decoy on the surface of filamentous particles. We find that the SEP-HA virion population contains ∼75% the amount of wildtype HA found in normal virions. At steady-state, both S139/1 and C05 IgG binding remains proportional to the HA content, at ∼75% for SEP-HA virions relative to WT (Fig. 3B). This indicates that the decrease in antibody binding is likely the direct result of the decrease in available HA, rather than a change in bivalent binding; thus, a 25% decrease in surface density is not sufficient to diminish the avidity of either S139/1 or C05 IgG.

### Bivalent binding of S139/1 and C05 persists after ∼10-fold reductions in HA surface densities

To probe the relationship between HA surface density and antibody binding more systematically, we used 200 nm streptavidin beads, which we can decorate with C-terminally biotinylated HA at different surface densities. Briefly, the beads are immobilized onto a coverslip, and purified HA from the HK68 strain that is fluorescently-labeled and biotinylated via a C-terminal ybbR tag^26^ is introduced and allowed to bind to saturation. After washing to remove unbound HA, fluorescently-labeled antibody is added and allowed to reach equilibrium. To vary the HA surface density, biotinylated HA is mixed with varying concentrations of biotinylated BSA as a competitor (Fig. 3C). In this way, we can measure steady-state antibody binding over a wide range of HA surface densities, with the highest density (100%) most closely mimicking a wild-type viral surface. We confirmed that surface densities of HAs on the beads are similar to those found on viruses by comparing binding of fluorescent C05 Fab to beads at 100% HA density relative to strain-matched virions. Accounting for differences in size^27^, we estimate that HA density on the nanoparticle surface is ∼85% that of the average HK68 virion (Fig S2D).

For both S139/1 and C05 Fab, binding increases linearly with HA density, as expected for a monovalent interaction dictated by absolute HA availability rather than density (Fig. 3D). Interestingly, the same relationship is observed for S139/1 IgG. The amount of IgG bound is directly proportional to the amount of HA available, with no apparent decrease in antibody occupancy as HA density is reduced. For C05, we find that the normalized occupancy of HA (defined as the normalized antibody intensity divided by the normalized HA intensity) decreases slightly as the HA density increases, indicating that C05 antibody binds more effectively at lower HA densities. For both antibodies, the ratio of bound IgG to HA remains approximately constant down to densities as low as ∼10%, the lowest we tested (Fig. 3E). In contrast to the IgG, Fab binding measured at twice the molar concentration of the IgG is nearly undetectable under these conditions, confirming the IgG binding is not occurring through monovalent interactions (Fig. S2E). The different binding behavior of C05 as compared to S139/1 IgG antibodies is particularly surprising, as it is opposite the trend we would expect if antigen density were the primary determinant of avidity.

### Tilting of HA about its membrane anchor contributes to C05 and S139/1 avidity

These findings suggest that antigen characteristics beyond surface density contribute to the efficiency of bivalent antibody binding. To better understand how structural features of antibody-HA complexes may contribute to avidity and dependence on antigen density, we used a structure-based model^28^ to predict the orientations of adjacent HA trimers that could support bivalent binding by S139/1 and C05. By sampling different conformations of the antibody Fab domains and determining the resulting HA orientations from known structures, we obtain distributions for the linear distance between the base of the two HAs, as well as the angle between them (Fig. 4A). This analysis suggests that while C05 binding prefers HA trimers that are further apart and oriented towards each other, S139/1 binding prefers trimers that are closer together and with an orientation that is closer to parallel. Of note, the preferred spacing for both antibodies are greater than the estimated distance (∼10 nm) between proximal HAs on the native viral surface^6^. This suggests that the IgGs may bypass adjacent HAs to bind to secondary or even tertiary HA neighbors. Expanding this analysis to other known antibody-HA structures suggests that C05 is not unique (Fig. S3A); for example, F045-092 also shows preference for a large inter-HA distance, and an even wider angle than C05, but has been shown to bind several orders of magnitude more strongly as IgG than as a Fab^29^.

**Figure 4.**
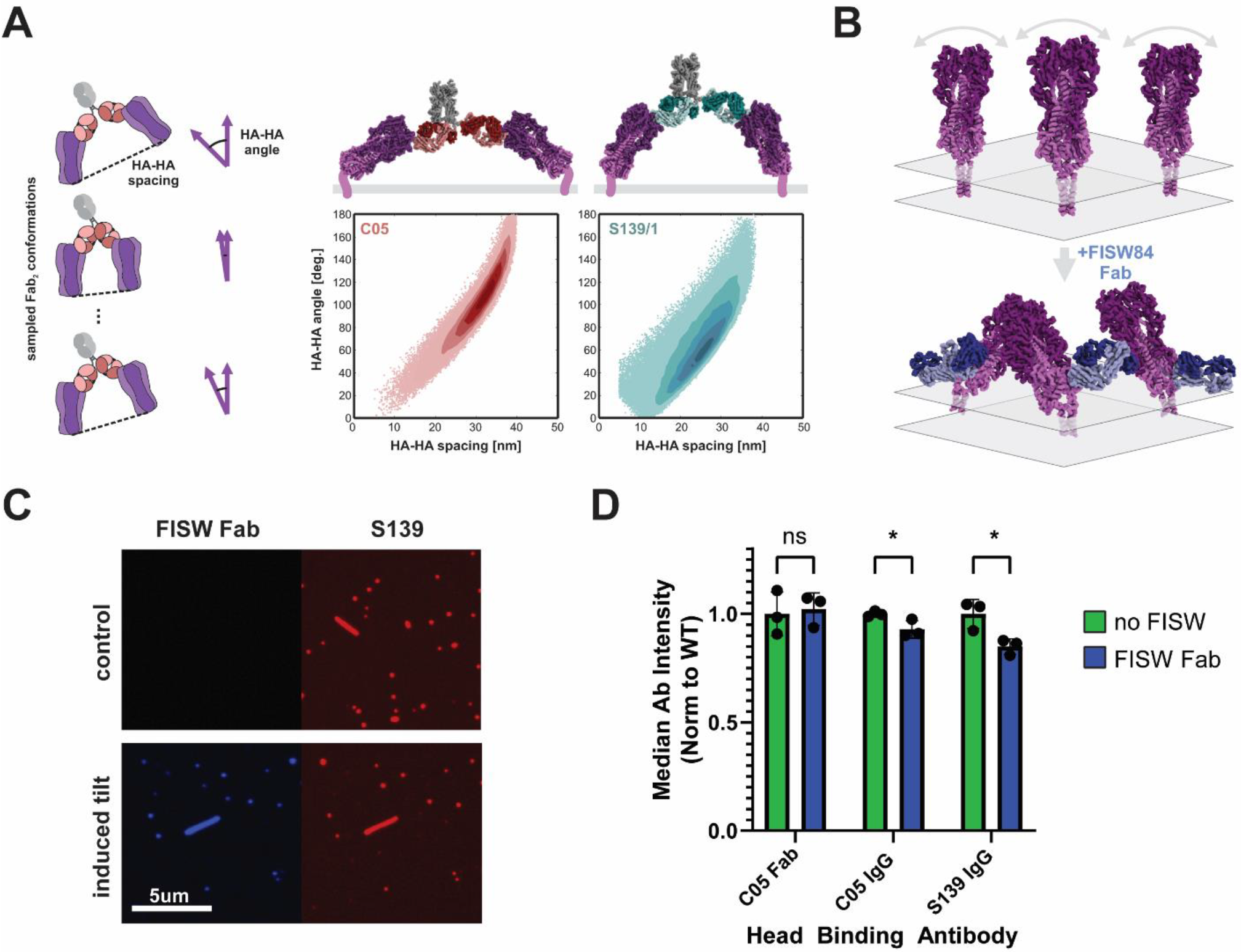
Antibodies prefer different HA orientations for efficient crosslinking. (A) Model to predict preferred antibody binding geometry. Structures of HA-Fab complexes are subjected to transformations reflecting the degrees of freedom of one Fab relative to the other. The distance between the base of the two bound HAs and the angle between them is determined for each sampled conformation. Plots in the lower right show angle and spacing distributions predicted for S139/1 and C05. Structural models above the plots are built by aligning HA-Fab structures (PDB IDs 4GMS and 4FQR) with human IgG1 (PDB ID 1HZH). (B) Schematic illustrating the proposed effect upon binding of an FISW84 Fab to HA (PDB IDs 6HJP, 6HJR). (C) Images of WSN33 bound by S139/1, +/-FISW84 Fab at 30nM. (D) Steady-state binding for the indicated antibodies. Analysis is performed on filamentous virions to ensure high signal-to-noise, and the median for each population is normalized to that of the corresponding wild-type condition. Three biological replicates are performed for each condition and compared by multiple unpaired t-tests; significance is indicated by a p-value below 0.05.

The HA ectodomain is connected to the transmembrane domain by a linker that permits extensive ectodomain tilting^30^. We hypothesized that this conformational flexibility may contribute to enhanced avidity by accommodating the structural preferences of different antibodies. If this is correct, we reasoned that avidity could be reduced by constraining tilting of the HA ectodomain. To test this hypothesis, we used FISW84, an antibody that binds to the HA anchor epitope and biases the ectodomain into a tilted conformation^30^ (Fig. 4B). Consistent with this hypothesis, viruses that are pre-bound by FISW84 Fab show reduced binding by S139/1 and C05 IgG, but not the C05 Fab (the affinity of the S139/1 Fab for A/WSN/1933 HA is not sufficiently strong for us to compare binding +/-FISW84) (Fig. 4C & D). Thus, although the FISW84 Fab recognizes an epitope >10nm away from that recognized by either C05 or S139/1, it is able to modestly inhibit the binding of these antibodies by perturbing the orientational dynamics or distributions of HA on the viral surface.

## DISCUSSION

While the importance of antibody avidity in virus binding and neutralization is well established^17,28,31–33^, the features of an antigenic surface that contribute to increased or decreased antibody avidity are not well understood. Using experimental approaches which directly perturb HA surface density and orientation, we are able to gain some insight into how these variables affect the avidity of two antibodies that bind to the HA head at different angles of approach – S139/1 and C05. We find that both antibodies are insensitive to at least a 10-fold decrease in HA surface density. This result is supported by structural predictions that suggest that these and other antibodies which bind to the HA head may rarely crosslink nearest neighbors within the viral membrane, binding instead to HAs whose anchors are separated by 20-30nm. It is important to note that we have focused here on IgG1 antibodies; differences in the hinge and valency of other antibody isotypes or subclasses are likely to be important^34,35^.

In addition to HA density, our data and analysis suggest that the ability of HA to tilt about its membrane anchor may play an important role in supporting bivalent attachment by some antibodies. While we predict that S139/1 can bridge two HAs tilted ∼30° towards each other with respect to the axis normal to the plane of the membrane, C05 favors an angle nearly twice as high (∼60°). In comparison, neighboring HA trimers on the viral surface would be expected to splay away from each other at an angle of ∼15° if they are oriented normal to the membrane^6^. This difference in preferred HA orientation may explain why S139/1 is more strongly affected by the presence of FISW84 (Fig. 4), which we expect will bias HAs into a more tilted conformation. It may also explain why C05 shows higher binding occupancy at low HA densities in experiments with synthetic nanoparticles (Fig. 3); lower HA densities may facilitate sampling of extreme angles that are obstructed when HA densities are high. Direct measurements of HA conformational dynamics in the presence or absence of antibodies will be needed to determine if this is the case.

The density of glycoproteins on the viral surface poses a potential evolutionary tradeoff; while high densities facilitate viral attachment to the cell surface, they may also make the virus more susceptible to neutralization by high-avidity antibodies. There is some evidence that different viruses have arrived at different evolutionary solutions to this problem. HIV-1 virions have only a few copies of the Env protein per virus – sufficient for binding the viral receptor through high affinity, but at a distance that is too far apart for an IgG to bind bivalently^36,37^. Without enhanced binding through avidity, neutralization must be achieved primarily through antibodies with a high monovalent affinity^31,38^. Influenza is at the opposite extreme, with neighboring glycoproteins separated by ∼10nm from each other. Our results suggests that reducing HA surface densities by up to 10-fold would have minimal effect on the avidity of the antibodies tested here. If this is the case, there would be little evolutionary pressure towards lower HA surface densities, which could reduce viral attachment in cases where high-affinity sialic acid receptors are limited without reducing the occupancy of some avidity-dependent antibodies. Besides HA density, our results and analysis suggest an important role for HA flexibility in increasing antibody avidity. This, too, could be important for viral fitness, for example, by allowing the HA ectodomain to pivot about its membrane anchor during the foldback step of membrane fusion, or to better engage cellular receptors. Genetic approaches to tune HA flexibility and incorporation into budding virions beyond what we demonstrate here will be critical in testing these predictions.

## MATERIALS AND METHODS

### Cell and virus cultivation

MDCK-II and HEK-293T cell lines used in the study were purchased from ATCC as authenticated cell lines (STR profiling). They were grown under standard conditions (37°C and 5% CO2) and cultured with cell growth medium containing Dulbecco’s modified Eagle’s medium (DMEM; Gibco), 10% fetal bovine serum (FBS; Gibco) and 1× antibiotic-antimycotic (Corning).

Standard reverse genetics techniques we used for virus rescue of the WSN33 and HK68 strains^39^. Briefly, co-cultures of HEK-293T and MDCK-II were transfected with plasmids containing each of the eight vRNA segments flanked by bidirectional promoters. The virus was harvested approximately 48 hours after transfection and plaque purified. For virus expansion, the plaques were passaged at a low MOI (∼0.001) in MDCK-II cells, in a virus growth medium containing Opti-MEM (Gibco), 2.5 mg/mL bovine serum albumin (Sigma-Aldrich), 1 μg/mL L-(tosylamido-2-phenyl ethyl) chloromethyl ketone (TPCK)-treated trypsin (Thermo Scientific Pierce), and 1× antibiotic-antimycotic (Corning). The viral stocks were further expanded for experiments by passaging at low MOI in MDCK-II cells. Here, a version of WSN33 with filamentous morphology was used. To obtain this phenotype, the WSN M1 sequence was replaced by that of A/Udorn/1972, as described previously^40^.

### Protein purification and labeling

Sequences for the VH and VL regions of the antibodies of interest were obtained from deposited sequences and cloned into expression vectors to make either full-length human IgG1 antibodies or Fab fragments. The heavy chain sequences were modified with a C-terminal ybbR tag for enzymatic labeling and purified as described^41^. Briefly, full-length antibodies were purified using protein A agarose beads (Thermo Scientific Pierce), while Fab fragments were affinity purified from a His6 tag using Ni-NTA Agarose Beads (Thermo Scientific Pierce). Adherent HEK293T cells were allowed to grow to ∼90% confluency, transfected with the heavy and light chain plasmids, and allowed to express for 7 days. During this time, the cells were cultured in Opti-MEM with 1× antibiotic-antimycotic and 2% FBS for Fab expression, and without FBS for IgG expression.

Recombinant HA for experiments with beads was produced by transfecting HEK293T cells with a pCAGGS expression vector containing the sequence for the HK68 HA ectodomain, followed by a foldon, His6-tag for affinity purification, and ybbR tag for enzymatic biotinylation using Sfp synthase^26^. After expressing for five days, HA was purified from the supernatant with Ni-NTA Agarose Beads (Thermo Scientific Pierce), and enzymatically biotinylated overnight at 4°C with Sfp. The HA was then chemically labeled at a ratio of approximately one dye molecule per HA trimer using Sulfo-Cy5 NHS dye (Lumiprobe). Sequences of constructs used in this work are given in Supplementary Information.

### Imaging flow chamber construction and functionalization

Imaging flow chambers were built with channels of 1mm width and 0.17mm height. The chambers were constructed by securing no. 1.5 thickness coverslips to acrylic covers with a double-sided adhesive (3M). The 1/16” acrylic backing was laser cut to form wells, and the adhesive was vinyl cut to shape the channels.

The coverslips were functionalized via a series of incubation steps carried out at room temperature. First, 90 μg/mL of BSA-biotin was flowed into the chambers and allowed to adsorb to the coverslip for two hours. Remaining BSA-biotin was washed with an excess of PBS, then incubated with 25 μg/mL streptavidin (Invitrogen) in PBS for 1 hour. Flow chambers were washed again with excess PBS and subsequently incubated with 25 μg/mL biotinylated *Erythrina cristagalli* lectin (Vector Laboratories) for 2 hours to capture viruses on the coverslip. The channels were washed a final time with PBS before introducing virus for imaging.

### Antibody dissociation assay

Virions were immobilized in the functionalized flow chambers. The fluorescently labeled antibody of interest was washed in, and binding was allowed to reach equilibrium. The imaging chamber was then set up on a Nikon TI2 confocal microscopy system, using a 60×, 1.40-NA objective. The antibody was washed out with PBS, and dissociation was observed through loss of fluorescence signal. Images were acquired in a timelapse, with frame rate set relative to the expected dissociation rate.

Antibodies used for dissociation measurements were labeled with Alexa Fluor 555, selected for its brightness and excellent photostability. To confirm that the loss of signal in dissociation measurements was not due to photobleaching, S139/1 IgG bound at steady state was imaged without washing. Sixty images were taken at a rate of 1s under the same acquisition settings, to ensure that the number of frames collected were in excess of the dissociation acquisitions while minimizing antibody exchange.

### SEP-HA cells lines and virus

To create the SEP-HA expressing cell line, lentivirus was produced by transfecting the packaging and envelope vectors pCMV8.91 and pMD2.G, along with transfer vector pHR-SIN into HEK29T cells. The SEP-HA construct consisted of SEP, a GCN4-pII trimerization domain, and the WSN33 HA transmembrane domain/cytoplasmic tail, followed by an internal ribosome entry site (IRES) for expression of puroR. MDCK-II cells were infected with this lentivirus and selected with puromycin to create a cell line that constitutively expresses SEP-HA. To achieve uniform expression of SEP-HA, we performed a clonal expansion from single cells in 96-well plates and selected the highest expressing clone (as determined by cell surface SEP signal) for subsequent experiments.

SEP-HA-containing influenza virus was obtained by infecting this cell line with HK68 at an MOI of ∼1, while wild type virus was collected from the parental MDCK cell line for comparison. Virus was collected 18 hours post infection and immobilized in flow chambers. Fluorescent antibody was washed into the channels and allowed to bind to equilibrium. Signal of both the SEP and antibody fluorophore were collected at steady-state following incubation for 3-4 hours. Filamentous particles were segmented during image processing to ensure the best signal-to-noise in data analysis. The mean intensity per pixel was measured for each filamentous virion (defined as a particle with a major axis at least four times as long as the minor axis).

### Synthetic nanoparticles

C-terminally biotinylated HA was bound to streptavidin-coated beads of 200nm diameter (Bangs Laboratories) to mimic the viral surface. A glass-bottom 96 well plate was cleaned by incubating wells with 3M NaOH for 30 minutes and washing thoroughly with PBS. The plate was then incubated with BSA-biotin at room temperature for 1 hour to functionalize the surface. The streptavidin beads were diluted into PBS with 0.05% tween (PBST) and added to the wells for 15 minutes.

Relative HA density on the surface of beads was titrated by mixing biotinylated HA with varying concentrations of biotinylated BSA. The 100% HA solution contained 16nM of the purified HA, while the 0% HA solution contained 150nM BSA-biotin; mixtures of these solutions were prepared to achieve the intermediate concentrations shown in Fig. 3. Each mixture was added to a well and allowed to bind to the beads at 4°C overnight. The excess protein was washed thoroughly with PBS, and fluorescently labeled antibodies were added to the wells and allowed to bind for 4 hours. Images were collected using a Nikon TI2 confocal microscope with a 60×, 1.40-NA objective. Relative HA density and antibody binding were determined by normalizing to HA fluorescence and antibody fluorescence intensity respectively in the 100% HA condition.

### Competition assay with FISW84

WSN33 virus was produced by infecting MDCK cells at an MOI of ∼1 and collecting the virus-containing supernatant at 18h post infection. Virus was immobilized into functionalized flow chambers. To induce HA tilt, a fluorescent Fab derived from the antibody FISW84 was added at 30nM and allowed to bind for 1 hour; the Fab form was used so as not to confound results with any potential steric hindrance from the Fc. The fluorescent antibody of interest was then added (along with 30nM of FISW84 for the tilt-induced condition). As before, filamentous particles were segmented during image processing to ensure high signal-to-noise in data analysis.

### Determining antibody crosslinking rates from experimental data

For the crosslinking model shown schematically in Fig. 2A, the evolution of singly-bound (*A*_1_) and doubly-bound antibodies (*A*_2_) over time is given by the equations:

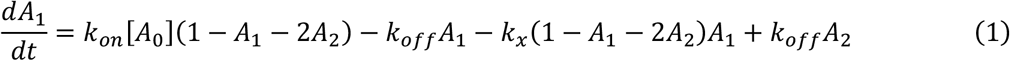

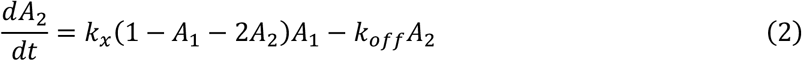

Here, *A*_1_ and *A*_2_ are normalized by the total number of HAs in the system, [*A*_0_] denotes the molar concentration of IgG in solution (assumed constant), *k*_*on*_ denotes the association rate for the IgG antibody (taken as twice the association rate of the Fab), *k*_*off*_ denotes the monovalent (Fab) dissociation rate, and *k*_*x*_ denotes the crosslinking rate. To determine values for *k*_*x*_ from measured IgG and Fab dissociation rates, we numerically integrate *A*_1_ and *A*_2_ and fit the sum (corresponding to the total bound antibody, the quantity that is captured by our measurements) to a kinetic model without crosslinking:

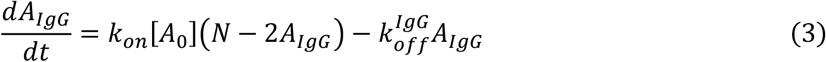

Here, 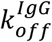 corresponds to the apparent dissociation rate measured for IgG antibodies (Fig. 2D), and we assume that all antibodies attach bivalently. For the high-avidity antibodies S139/1 and C05, this assumption is reasonable; for the concentrations of antibodies we use, our kinetic parameters suggest that *A*_2_ is typically ∼10-100-fold greater than *A*_1_. For fitting the crosslinking rate *k*_*x*_, we focus only on the dissociation phase, initializing the system using *A*_2_(*t* = 0) = *A*IgG(*t* = 0) = 0.2 and *A*_1_(*t* = 0) = 0. This fitting procedure is used to determine the values for *k*_*x*_ reported in Fig. 2D. It is important to note that as the initial value of *A*_2_ approaches its maximum value of 0.5, fitting becomes poor. Under these conditions, as the first doubly-bound antibodies dissociate into singly-bound antibodies, they are unable to efficiently re-form crosslinks because the majority of HAs remain occupied. This leads to an initial rapid dissociation phase which we do not observe in our experiments. This is a limitation of the continuum model, which does not preserve spatial information regarding the location of free HAs relative to antibodies that had previously been bound to them.

Solving (1) and (2) for the total fraction of bound HAs gives:

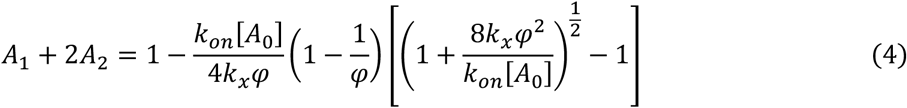

where

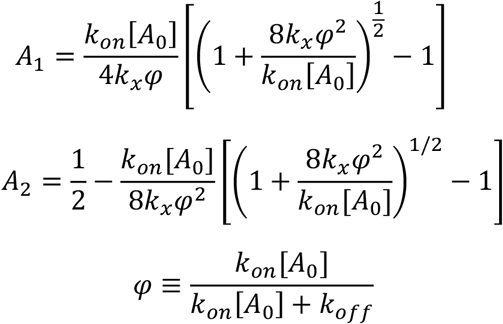

Setting (4) equal to ½ and solving for the antibody concentration [*A*_0_] allows us to determine the fold decrease in *K*_*D*_ for an IgG antibody relative to the Fab as a function of the crosslinking rate kx and other kinetic parameters. These results are plotted in Fig. S2. Matlab code for fitting and analysis is available on Github (https://github.com/mvahey/benegal2024).

### Statistics, replicates, and software

Statistical analysis was performed using GraphPad Prism 10 and MATLAB. The statistical tests and the number of replicates used in specific cases are described in the figure legends. No statistical methods were used to predetermine sample size.

## Supporting information

Protein Sequences

## Notes

https://github.com/mvahey/benegal2024

### Competing Interest Statement

The authors have declared no competing interest.

