## Supplemental Figures for "Antigen flexibility supports the avidity of hemagglutinin-specific antibodies at low antigen densities"

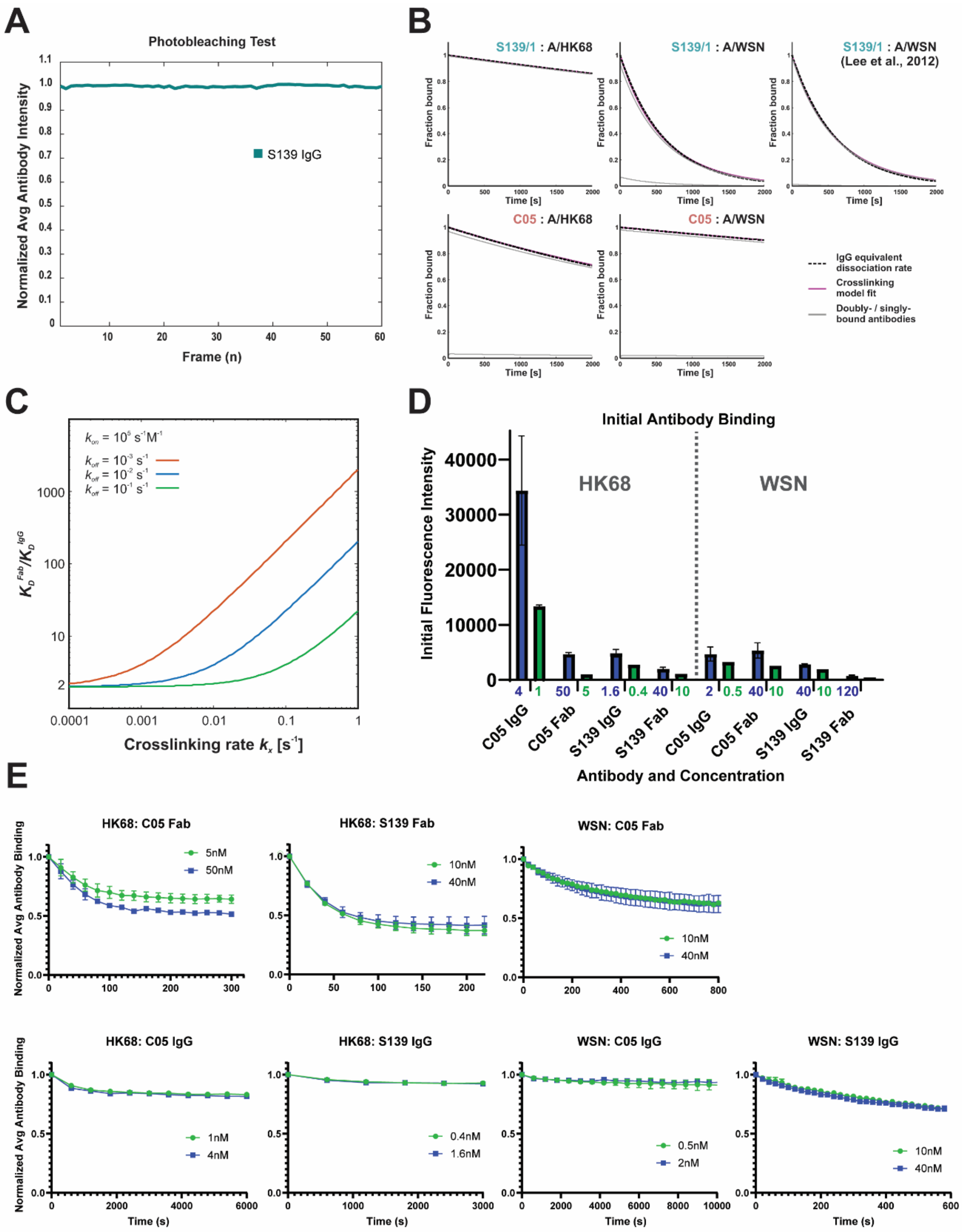


**Figure S1. Characterization of antibody binding and crosslinking kinetics.** (A) To isolate the effects of photobleaching, fluorescent antibody was bound to immobilized virus and imaged with the same acquisition settings as the dissociation experiments, but without washing out the antibody. The signal is not significantly changed over 60 time frames. (B) To fit the crosslinking rate ($k_x$), the experimentally obtained $k_{off}$ values from each Fab and IgG pair were mapped to the simulation results. For a given Fab $k_{off}$ value, we iterate through a series of $k_x$ values and determine the best fit to the dissociation curve for the corresponding IgG. (C) The fold difference in effective $k_{off}$ is shown as a function of $k_x$ for a few given Fab $k_{off}$ values. (D) Intensities of bound antibodies prior to measuring dissociation. Data is from the experiments shown normalized in Figure 1C. (E) Normalized dissociation curves for the conditions plotted in D.

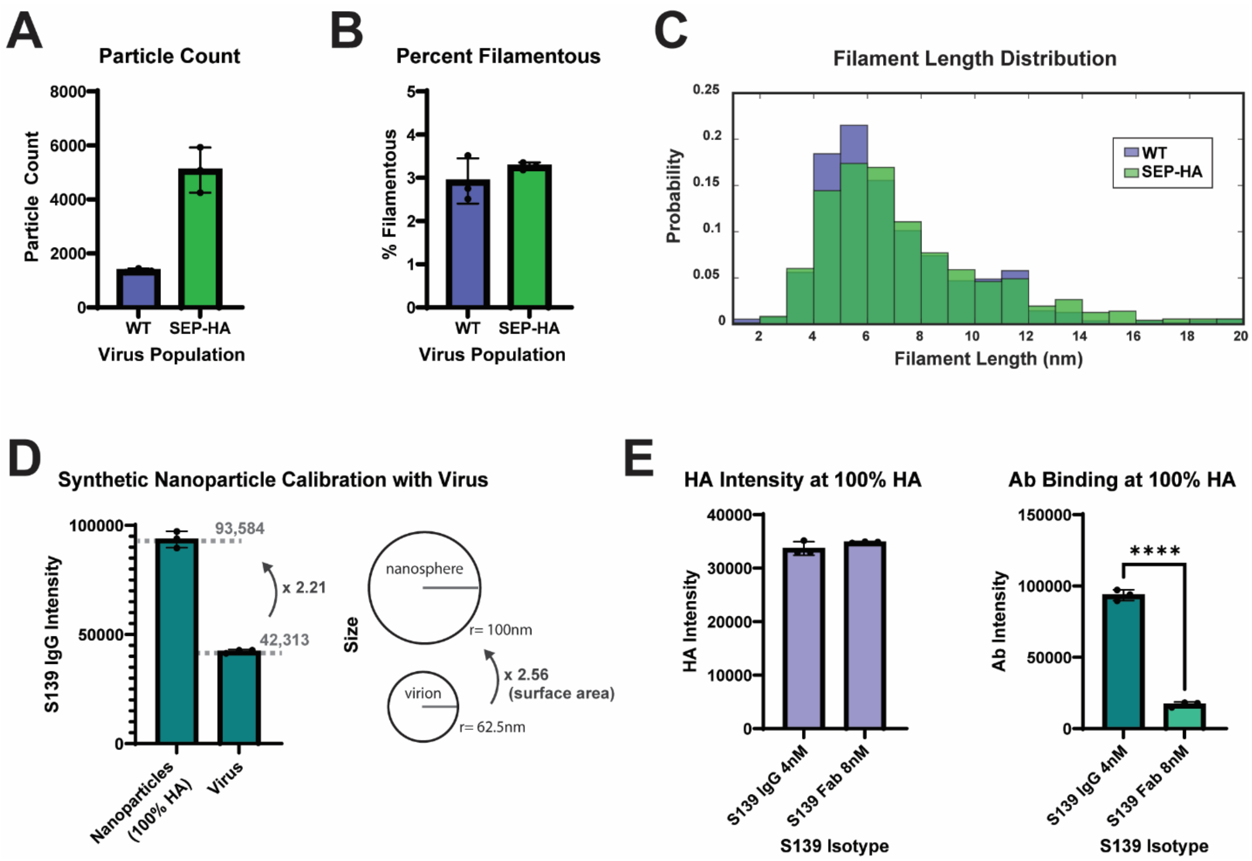


**Figure S2. Determining the effects of reduced antigen density on antibody avidity.** Comparisons of wildtype and SEP-HA virus populations based upon (A) total particle count; (B) percentage filamentous particles; and (C) filament length distributions. (D) Calibration of HA density on streptavidin nanoparticles relative to native virions. Plot to the left quantifies the fluorescent intensity of labeled S139/1 IgG bound to nanoparticles (incubated with biotinylated HA) and viruses. Schematic to the right shows relative sizes of nanoparticles and virions used to calculate relative HA densities. The corresponding scaling between the size of an average spherical virion to a nanoparticle is shown. (E) Quantification of HA intensities (left plot) and antibody intensities (right plot) for beads with saturating densities ('100%') of HA, incubated with S139/1 IgG or S139/1 Fab.

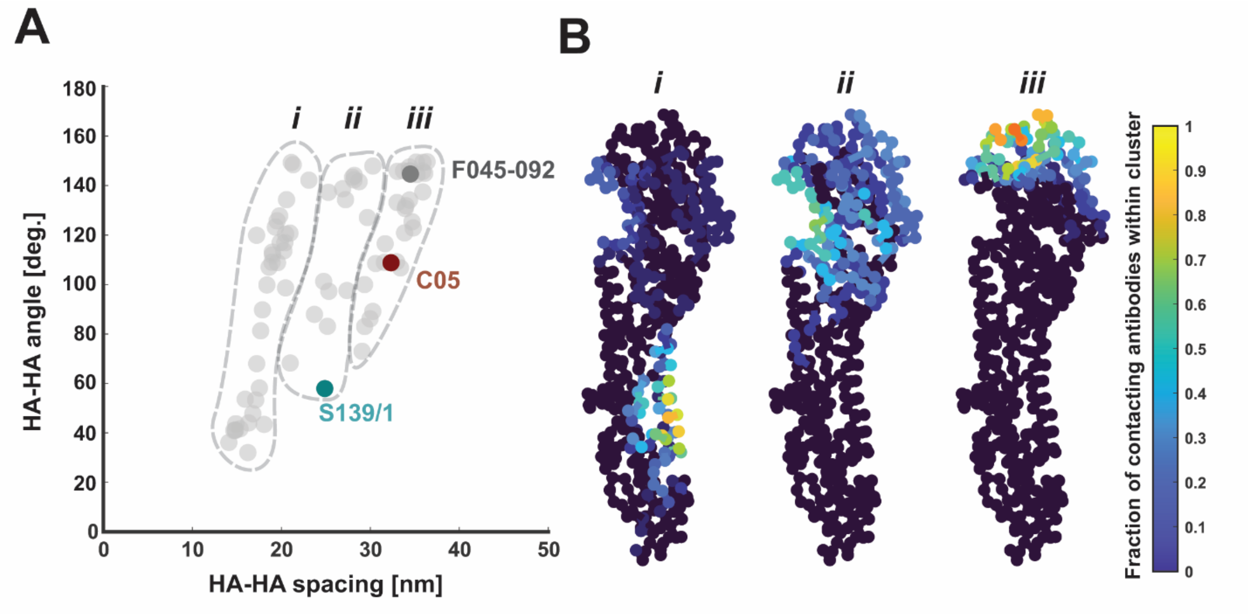


**Figure S3. Modeling the geometric preferences of HA-specific monoclonal antibodies.** (A) Predictions from the structure-based model from Fig. 4A, extended to structures of HA-antibody complexes from the PDB. Each point represents the most frequent inter-HA spacing and angle samples by a particular antibody. S139/1, C05, and F045-092, another high-avidity antibody, are highlighted. (B) Structure of an HA monomer with each residue colored according to the frequency of contacts by antibodies in the corresponding groups from the plot in *A*. Contacts are defined as HA residues within 0.8 nm of any residue from the antibody heavy or light chain.
