## Supplementary material for "Antigen flexibility supports the avidity of hemagglutinin-specific antibodies at low antigen densities": Protein Sequences

S139/1 HC:
MGWSCIILFLVATATGVHSEVQLQQSGTELKKPGASVKISCKATGYTFSSYWIEWIKQRPGHGLEWIGEILPEIGMTNYNENFKGKATFTANTSSNTVY
MQLSSLTSEDSAVYYCARPYDYSWFAYWGQGTLVTVSSASTKGPSVFPLAPSSKSTSGGTAALGCLVKDYFPEPVTVSWNSGALTSGVHTFPAVLQSSG
LYSLSSVVTVPSSSLGTQTYICNVNHKPSNTKVDKRVEPKSCDKTHTCPPCPAPELLGGPSVFLFPPKPKDTLMISRTPEVTCVVVDVSHEDPEVKFNW
YVDGVEVHNAKTKPREEQYNSTYRVVSVLTVLHQDWLNGKEYKCKVSNKALPAPIEKTISKAKGQPREPQVYTLPPSRDELTKNQVSLTCLVKGFYPSD
IAVEWESNGQPENNYKTTPPVLDSDGSFFLYSKLTVDKSRWQQGNVFSCSVMHEALHNHYTQKSLSLSPGK*

S139/1 HC Fab:
MGWSCIILFLVATATGVHSEVQLQQSGTELKKPGASVKISCKATGYTFSSYWIEWIKQRPGHGLEWIGEILPEIGMTNYNENFKGKATFTANTSSNTVY
MQLSSLTSEDSAVYYCARPYDYSWFAYWGQGTLVTVSSASTKGPSVFPLAPSSKSTSGGTAALGCLVKDYFPEPVTVSWNSGALTSGVHTFPAVLQSSG
LYSLSSVVTVPSSSLGTQTYICNVNHKPSNTKVDKRVEPKSCDKGGSHHHHHHGGSDSLEFIASKLA*

S139/1 LC:
MGWSCIILFLVATATGVHSDIVMTQSQKFMSTSVGDRVSVTCKASQNVDTNVAWYQEKPGQSPKTLIYSASNRYSGVPDRFTGSASGTDFTLTITNVQS
EDLAEYFCQQYNSYPYTFGGGTKLEIKRADAAPSVFIFPPSDEQLKSGTASVVCLLNNFYPREAKVQWKVDNALQSGNSQESVTEQDSKDSTYSLSSTL
TLSKADYEKHKVYACEVTHQGLSSPVTKSFNRGEC*

C05 HC:
MGWSCIILFLVATATGVHSEVQLQESGGGLVQPGESLRLSCVGSGSSFGESTLSYYAVSWVRQAPGKGLEWLSIINAGGGDIDYADSVEGRFTISRDNS
KETLYLQMTNLRVEDTGVYYCAKHMSMQQVVSAGWERADLVGDAFDVWGQGTMVTVSSASTKGPSVFPLAPSSKSTSGGTAALGCLVKDYFPEPVTVSW
NSGALTSGVHTFPAVLQSSGLYSLSSVVTVPSSSLGTQTYICNVNHKPSNTKVDKRVEPKSCDKTHTCPPCPAPELLGGPSVFLFPPKPKDTLMISRTP
EVTCVVVDVSHEDPEVKFNWYVDGVEVHNAKTKPREEQYNSTYRVVSVLTVLHQDWLNGKEYKCKVSNKALPAPIEKTISKAKGQPREPQVYTLPPSRD
ELTKNQVSLTCLVKGFYPSDIAVEWESNGQPENNYKTTPPVLDSDGSFFLYSKLTVDKSRWQQGNVFSCSVMHEALHNHYTQKSLSLSPGK*

C05 HC Fab:
MGWSCIILFLVATATGVHSEVQLQESGGGLVQPGESLRLSCVGSGSSFGESTLSYYAVSWVRQAPGKGLEWLSIINAGGGDIDYADSVEGRFTISRDNS
KETLYLQMTNLRVEDTGVYYCAKHMSMQQVVSAGWERADLVGDAFDVWGQGTMVTVSSASTKGPSVFPLAPSSKSTSGGTAALGCLVKDYFPEPVTVSW
NSGALTSGVHTFPAVLQSSGLYSLSSVVTVPSSSLGTQTYICNVNHKPSNTKVDKRVEPKSCDKGGSHHHHHHGGSDSLEFIASKLA*

C05 LC:
MGWSCIILFLVATATGVHSDIQLTQSPSSLSASVGDRVTLTCQASQDIRKFLNWYQQKPGKGPKLLIYDASNLQRGVPSRFSGGGSGTDFTLIISSLQP
EDVGTYYCQQYDGLPFTFGGGTKVVIKRTVAAPSVFIFPPSDEQLKSGTASVVCLLNNFYPREAKVQWKVDNALQSGNSQESVTEQDSKDSTYSLSSTL
TLSKADYEKHKVYACEVTHQGLSSPVTKSFNRGEC*

FISW84 Fab HC:
MGWSCIILFLVATATGVHSEVQLLESGGGLVQPGGSLRLSCAASGFTFSSYGMAWVRQAPGKGLEWVSFISATGLSTYFADSVKGRFTISRDTTKNTLY
LQMNSLRADDTAVYFCARMRRTMIAFGGNDFWGQGTLVTVSSASTKGPSVFPLAPSSKSTSGGTAALGCLVKDYFPEPVTVSWNSGALTSGVHTFPAVL
QSSGLYSLSSVVTVPSSSLGTQTYICNVNHKPSNTKVDKRVEPKSCDKGGSHHHHHHGGSDSLEFIASKLA*

FISW84 LC:
MGWSCIILFLVATATGVHSEVVMTQSPATLSVSPGEGATLSCRASQSVNTNVAWYQQKPGQAPRLLIYGASTRATGIPARFSGSGSGTEFTLTISTLQS
EDFAVYYCQQYSNWPPITFGQGTRLEIKRTVAAPSVFIFPPSDEQLKSGTASVVCLLNNFYPREAKVQWKVDNALQSGNSQESVTEQDSKDSTYSLSST
LTLSKADYEKHKVYACEVTHQGLSSPVTKSFNRGEC*

A/Hong Kong/1968 HA ectodomain:
MKTIIALSYIFCLALGQDLPGNDNSTATLCLGHHAVPNGTLVKTITDDQIEVTNATELVQSSSTGKICNNPHRILDGIDCSLIDALLGDPHCDVFRNET
WDLFVERSKAFSNCYPYDVPDYASLRSLVASSGTLEFITEGFTWTGVTQNGGSNACKRGPGSGFFSRLNWLTKSGSTYPVLNVTMPNNDNFDKLYIWGV
HHPSTNQEQTSLYVQASGRVTVSTRRSQQTIIPNIGSRPWVRGLSSRISIYWTIVKPGDVLVINSNGNPIAPRGYFKMRTGKSSIMRSDAPIDTCISEC
ITPNGSIPNDKPFQNVNKITYGACPKYVKQNTLKLATGMRNVPEKQTRGLFGAIAGFIENGWEGMIDGWYGFRHQNSEGTGQAADLKSTQAAIDQINGK
LNRVIEKTNEKFHQIEKEFSEVEGRIQDLEKYVEDTKIDLWSYNAELLVALENQHTIDLTDSEMNKLFEKTRRQLRENAEDMGNGCFKIYHKCDNACIE
SIRNGTYDHDVYRDEALNNRFQIKGVELKSGYKDGYIPEAPRDGQAYVRKDGEWVLLSTFLGSGSHHHHHHGGDSLEFIASKLA*

SEP-HA (A/WSN/1933):
MVEMLPTVAVLVLAVSVVAKDNTTLQEFATMVKGEELFTGVVPILVELDGDVNGHKFSVSGEGEGDATYGKLTLKFICTTGKLPVPWPTLVTTLTYGVQ
CFSRYPDHMKRHDFFKSAMPEGYVQERTIFFKDDGNYKTRAEVKFEGDTLVNRIELKGIDFKEDGNILGHKLEYNYNDHQVYIMADKQKNGIKANFKIR
HNIEDGGVQLADHYQQNTPIGDGPVLLPDNHYLFTTSTLSKDPNEKRDHMVLLEFVTAAGITHGMDELYKGGENLYFQGGGGSKQIEDKIEEILSKIYH
IENEIARIKKLIGGSGVYQILAIYSTVASSLVLLVSLGAISFWMCSNGSLQCRICI*
